# Fever temperatures modulate intraprotein dynamics and enhance the binding affinity between monoclonal antibodies and the Spike protein from SARS-CoV-2

**DOI:** 10.1101/2022.10.24.513610

**Authors:** Dong Gun Kim, Hak Sung Kim, Yoonjoo Choi, Razvan C. Stan

## Abstract

Fever is a typical symptom of most infectious diseases. While prolonged fever may be clinically undesirable, mild reversible fever (< 39°C, 312K) can potentiate the immune responses against pathogens. Here, using molecular dynamics, we investigated the effect of febrile temperatures (38°C to 40°C, 311K to 313K) on the immune complexes formed by the SARS-CoV-2 spike protein with two neutralizing antibodies. We found that, at mild fever temperatures (311-312K), the binding affinities of the two antibodies improve when compared to the physiological body temperature (37°C, 310K). Furthermore, only at 312K, antibodies exert distinct mechanical effects on the receptor binding domains of the spike protein that may hinder SARS-CoV-2 infectivity. Enhanced antibody binding affinity may thus be obtained using appropriate temperature conditions.

---

SARS-CoV-2 is a new coronavirus responsible for the ongoing pandemic of coronavirus disease 2019 (COVID-19). To gain entry into human cells, SARS-CoV-2 uses the transmembrane viral S glycoprotein (Spike) that protrudes from the viral surface and constitutes the main target of current vaccines [1]. On COVID-19 infection, fever is a main symptom and a key feature of the immune response [2]. Fever that accompanies COVID-19 is generally treated with the aim of eliminating it [3]. Recently, a study showed that at 313K (40°C), Spike attachment to ACE2 receptor is impaired [4]. It is unclear, however, what the role of fever may be on the formation of immune complexes. Here, we show that protein complexes formed between therapeutic antibodies and Spike proteins benefit from the effects of febrile temperatures. We chose two potent monoclonal antibodies (mAb), CV30 and S309, which are reported to neutralize SARS-CoV-2 [5, 6]. We indicate that at 312K (39°C), the binding affinity of mAb towards Spike is highest when compared to 310K (37°C, core body temperature), 311K (38°C, mild fever) or 313K (40°C, acute fever). Further, the detailed modes of the mechanisms of antibody neutralization under fever temperatures were uncovered: at 312K, CV30 “pulls away” the receptor binding domain (RBD_1_) off the Spike monomer it is bound to, from the RBD_2_ and RBD_3_ belonging to the other chains that do not contact the mAb. In so doing, CV30 exposes the RBD_2,3_ to cell receptor binding or to further mAb attachment. In contrast, at 312K and at 313K, the S309 has the reverse role on Spike organization and in effect closes the trimer structure, rendering cell entry impossible. An overview of the dynamics between Spikes and these mAb at relevant temperatures is shown in Figure 1.

**Figure 1.**
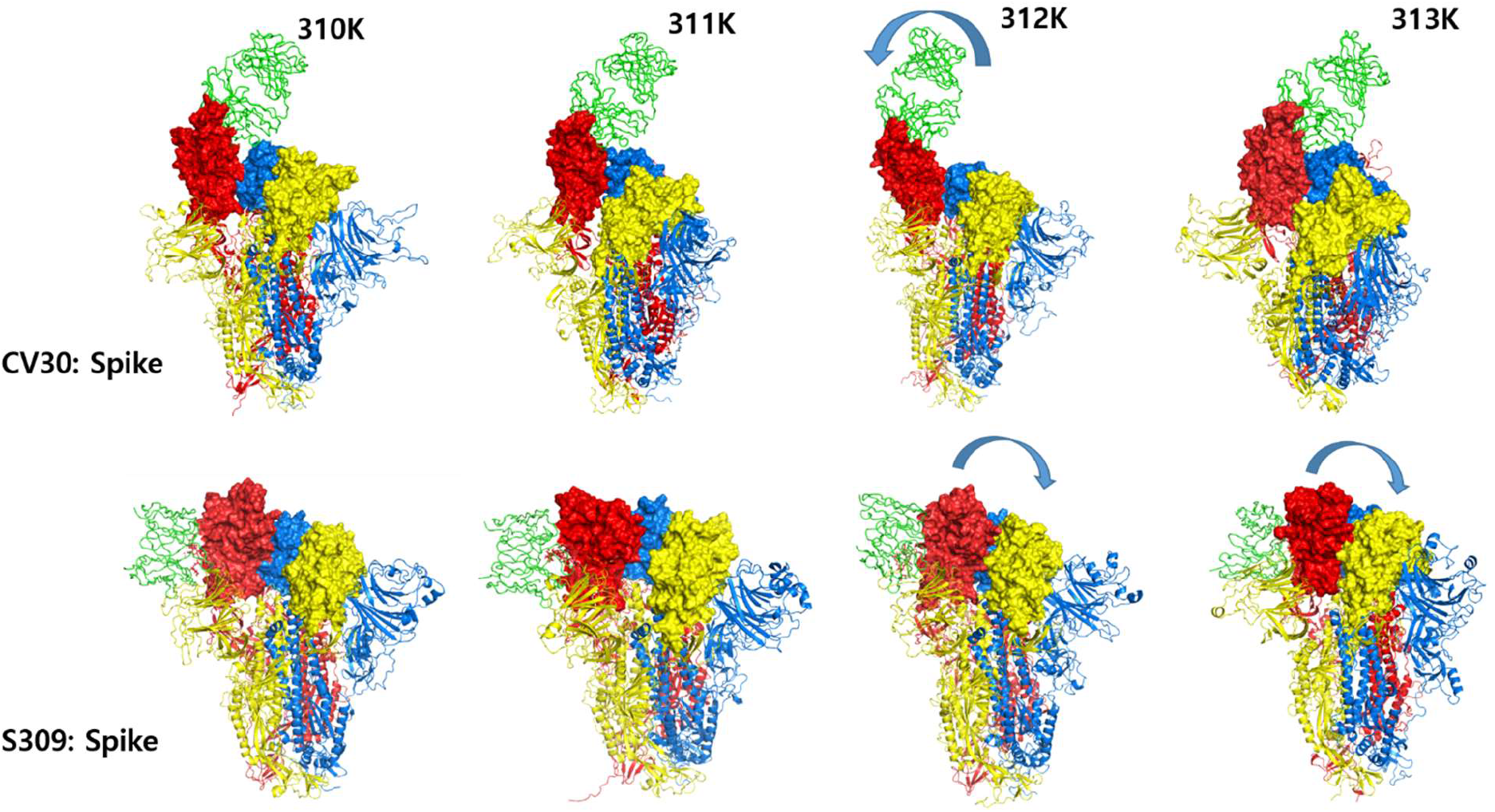
Snapshots taken after 500 ns of the Spike proteins in complexes with CV30 or with S309, at the indicated temperatures. Chain A (red cartoon) contains the RBD_1_ that is solely bound to respective antibodies (shown as green ribbons). Chains B and C are colored in yellow or blue cartoons, respectively; RBDs are shown as surfaces. Arrows indicate the translational movements of RBD_1_ at 312K or at 313K.

We established the center of mass (COM) for each of the three RBDs and determined the extent of translations with respect to each other, at fixed intervals of 0 ns, 250 ns and 500 ns, using the positions of COM of each RBD at 310K are used as references (Supplementary Table 1). For the CV30: Spike system, at 312K, the distances between RBD_1_ (chain A) versus RBD_2,3_ of chain B or chain C increase by 6.2Å and 20.1Å, respectively. For the S309: Spike complex, at 312 K, RBD_1_ approaches RBD_2_ by 10.6Å, while at 313K, the distance between same RBDs is reduced by 13.8Å. Further, to gauge the extent of the changes in position of each RBD with respect to itself, across all temperatures, we determined the shifts along their rotation axes, using the RBD positions at 310K as reference (Table 1).

**Table 1.** Translational shifts along the axes of rotations between the position of RBDs at 310 K and the positions of RBDs from the immune complexes under relevant febrile temperatures. The negative sign indicates translations away from the rotation axes.

| Shift along axis of rotation (Å) | CV30: Spike |  |  | S309: Spike |  |  |
| --- | --- | --- | --- | --- | --- | --- |
|  | 311K | 312K | 313K | 311K | 312K | 313K |
| RBD<sub>1</sub> (0 ns) | +2.1 | -1.1 | -1.5 | +0.2 | +0.1 | +0.2 |
| RBD<sub>1</sub> (500 ns) | +13.4 | -35 | -5.8 | +3.5 | +16.4 | +0.9 |
| RBD<sub>2</sub> (0 ns) | -1.2 | -0.3 | -0.4 | +0.2 | -0.2 | -0.04 |
| RBD<sub>2</sub> (500 ns) | +20 | -40 | -13 | +3.9 | +14 | +14 |
| RBD<sub>3</sub> (0 ns) | +1 | +0.6 | +1.1 | +0.2 | -0.1 | -0.3 |
| RBD<sub>3</sub> (500 ns) | +15 | -58 | +15 | +5.5 | +16 | +10.7 |

At 312K, each RBD in the CV30: Spike complex exhibits the largest translation away against own positions at 310K, while the opposite situation is encountered for the RBDs from the S309: Spike immune protein complex at 312K and 313K. For either immune complex, the origins of these RBD movements within the Spike trimer are found in the presence of short β-strands (RBD_1_ sequence residues 354-358, 452-455 and 492-494) that are connected by interstrands hydrogen bonds and help maintain a rigid binding interface with CV30 at 310, 311 and 313K. Conversely, at 312K, these secondary structures are lost and enhanced flexibility is gained that allows for the RBD_1_ swing motions away from the other two RBDs (Supplementary Figure 1). While the same phenomenon occurs with the immune complex formed with S309, the presence of β-strands (at 310K and 311K) or their absence (at 312K and 313K) involves residues that are not part of RBD_1_ (residues 312-319); additionally, more extended β-strands are comparatively found at 310K and 311K, that help maintain the position of RBD_1_ with respect to other RBDs (residues 592-594, 661-663).

We quantified the extent of global structural changes across temperatures using the RMSD and R*g* parameters (Supplementary Figure 2) as well as complex formation or dissociation between the mAb and the Spikes, using the Buried Surface Area (BSA) parameter (Table 2) and. For the CV30 bound to Spikes, BSA decreases by 18% at 312K, mirroring the movement away from the other RBDs, while it monotonically increases from 310K to 312K. In contrast, for the S309: Spike system, the increase in BSA is nonlinear with temperatures, peaking at 312K, in line with RBD_1_ movement towards the other RBDs.

**Table 2.** Changes in BSA between the mAb and the Spike proteins throughout the simulations at the indicated temperatures. In brackets, percentage increase or decrease against values at 0 ns.

| BSA<br>(Å<sup>2</sup>) | 310K CV30:<br>Spike | 311K<br>CV30: Spike | 312K<br>CV30: Spike | 313K<br>CV30: Spike | 310K S309:<br>Spike | 311K S309:<br>Spike | 312K<br>S309: Spike | 313K<br>S309: Spike |
| --- | --- | --- | --- | --- | --- | --- | --- | --- |
| 0 ns | 1285 | 1296 | 1533 | 1313 | 1287 | 1078 | 1068 | 1327 |
| 500<br>ns | 1412 (+9%) | 1480 (+14%) | 1260 (-18%) | 1679 (+27%) | 1534 (+19%) | 1510 (+40%) | 1660 (+55%) | 1936 (+45%) |

Next, we determine the extent of intra-monomer movements, using again BSA as a proxy for the formation of interfaces, as shown in Supplementary Table 2. While modest changes in BSA within the Spike monomers occur for the CV30: Spike complex (a 7% increase at 312K and at 313K with respect to 310K), a 21% peak increase at 312K versus 310K is present in the S309: Spike immune complex. Dissociation between the Spike monomers has two key roles for viral infectivity: to expose the fusion molecular machinery required for cell entry [7] and to promote RBD opening necessary for binding to cell entry receptor ACE2 [8]. Fever modulation of increased intra-monomer BSA may thus affect Spike stability, such that viral entry could be impaired.

In order to further quantify the strength in the binding affinity between the mAb and the RBD_1_, we estimate the changes in hydrogen bond formation across temperatures (Supplementary Figure 3 and Supplementary Table 4) and further calculated the free binding energy, as shown in Table 3.

**Table 3.** Interaction energies between Spike and the two paratopes, at physiological and at febrile temperatures. In brackets, percentage increase or decrease with respect to the reference values at 310K. Standard deviations are approximately 10% for the CV30: RBD_1_ complex and 15% for the S309:RBD_1_ system, respectively. Values taken from structures obtained after 500 ns of simulations.

| Binding Free<br>Energy (kCal/mol) | 310K | 311K | 312K | 313K |
| --- | --- | --- | --- | --- |
| CV30: RBD<sub>1</sub> | -18.6 | -24 (+29%) | -23 (+23%) | -19 (+2%) |
| S309: RBD<sub>1</sub> | -71 | -59 (-17%) | -96 (+35%) | -88 (+23%) |

In agreement with the results presented in Tables 1-3, we measured an inverse relation between increasing temperatures and the binding energy of CV30 in complex with RBD_1_, peaking at 311K. In contrast, the highest affinity is present at 312K for S309 bound to RBD_1_, in line with the movements of this RBD within the Spike at various febrile temperatures.

To control the ongoing pandemic, it is imperative to modulate the fundamental dynamics of the SARS-CoV-2 Spike, as its motions are key to the infection machinery [9]. Despite the increasing interest in the use of fever as an adjuvant to therapy [10], temperature modulation of immune complex formation, has just recently been put investigated, but only at 313K [4, 11]. Observational studies reported worse outcomes among COVID-19 patients having a lower body temperature (<36°C, 309K) on admission, suggesting an impaired immune response [12]. We here present the first results wherein a range of febrile temperatures are shown to: (1) modulate domain movements in a viral protein and (2) increase mAb binding affinity against viral epitopes at particular fever values. In this study, antibodies bind directly to RBD, as with CV30 [5], or away from RBD, as in the case of S309 [6], revealing a possible allosteric effect of febrile temperatures in the formation of immune complexes. Fever is a beneficial [13] and conserved [14] reaction to acute infections. Temporarily maintaining a fever temperature in relevant patients may thus potentiate the role of antibodies in recognizing viral epitopes and in reducing viral infectivity.

## Methods

The complex structure of SARS-CoV-2 spike and each antibody was obtained from the Protein Data Bank (PDB ID: 6VSB for Spike alone, 6WPS for S309 and 6XE1 for CV30). As the complex structure of CV30 was defined only using the receptor binding domain (RBD) of the spike protein, the spike bound state was modelled using Modeller. Simulations were performed using GROMACS 2020.2. Structures were pre-minimized using amber99sb forcefield with SPC/E explicit solvent model, equilibrated to each tested temperature (310-313 K) using NVT ensemble, followed by NPT ensemble equilibration. The production run was conducted for 500 nanoseconds (400 nanoseconds for Spike alone system) with 2 femtosecond time step and snapshots were taken every 10 picoseconds. The leapfrog integration method, V-rescale thermostat and Parrinello-Rahman barostat were used. LINCS constraints were applied to hydrogen bonds and long range interactions were included using Particle Mesh Ewald method. The changes in root-mean-square deviation (RMSD) of the whole complex, radius of gyration of the system, and the total solvent accessible surface area (SASA) vs. time were calculated using GROMACS. The preservation of the binding event was examined from the changes in BSA of receptor binding motif (Asn437-Tyr508). Binding free energy was computed using MM-PBSA method from designated snapshots. Center of mass for each RBD and RBD translational movements with respect to each other were calculated using Chimera and the Domain Rotation tool.

## Acknowledgments

We thank Dr. Aura Precupas for the initial input on this manuscript. This work was funded by the Korean National Supercomputing Center (grant KSC-2020-CRE-0203). Y.C. was supported by National Research Foundation of Korea (NRF-2020M3A9G3080281 and NRF-2020R1A5A2031185). RSC was supported by Chonnam National University (grant 2022-2574) and by National Research Foundation of Korea (NRF-2021R1I1A2059587).

## Supporting Information

Methodology; RBD movements within the Spike trimer; Dynamics of hydrogen bonds formation; Changes in RMSD and *R*g across febrile temperatures; Changes in the RBDs’ centers-of-mass with respect to each other; Changes in Buried Surface Areas between immune complexes and Kolmogorov–Smirnov tests for RMSD and Rg of all proteins at fever temperatures referenced against values at 310K.

## Author Contributions

RSC conceived of the study. RSC and YC secured funding. DGK performed the experiments. RSC wrote the manuscript. DGK, HSK, YC revised the final draft.

## Classification

Biophysics and computational biology.

---

**Supplementary Figure 1.**
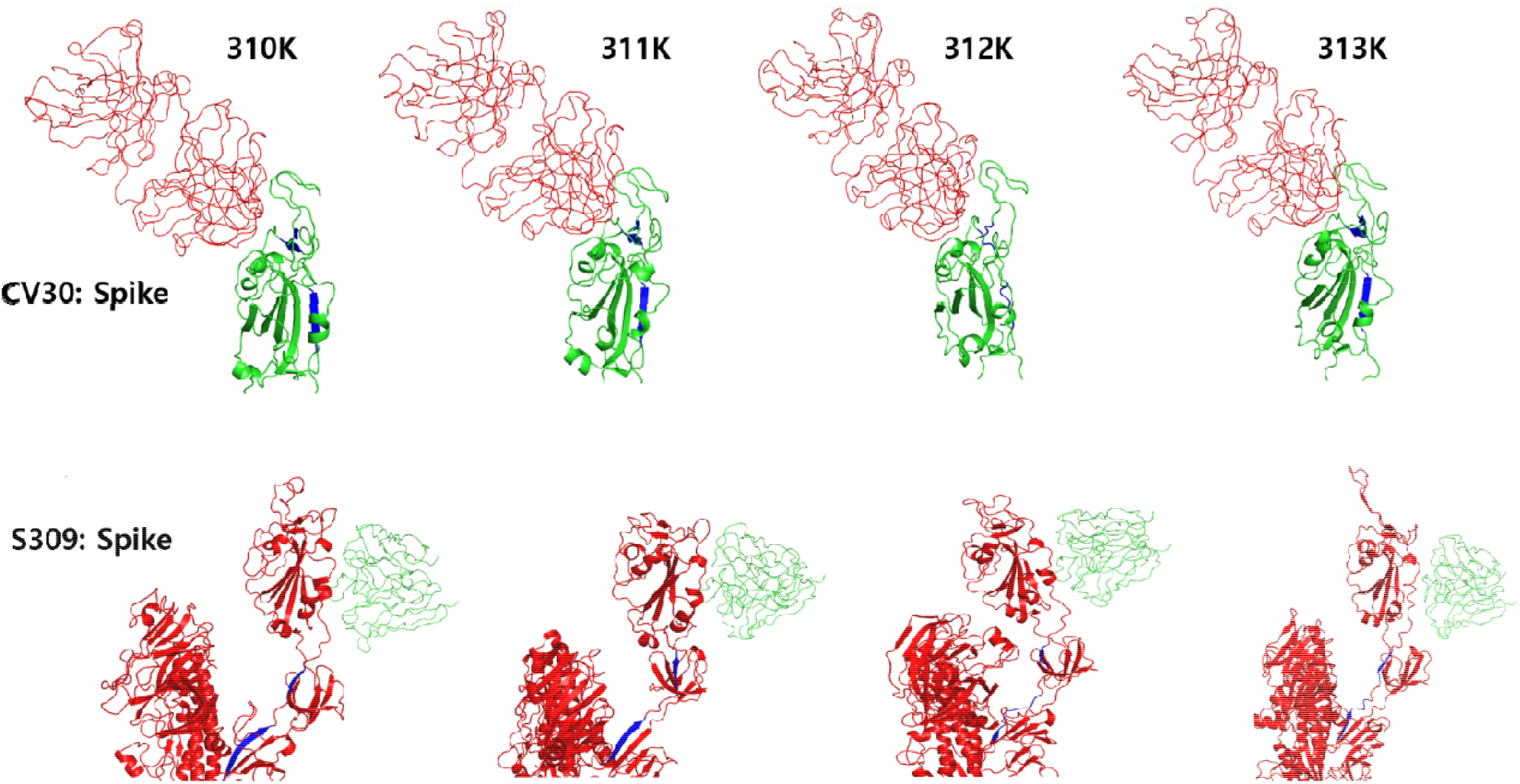
RBD_1_ swing motions away from the other two RBDs at different temperatures. Monoclonal antibodies are shown in red (top panel) or in green (bottom panel). In blue, the beta strands that are present at 310K, 311K and 313K but absent at 312K in both immune complexes.

We investigated the conformational transitions from the native to possible heat-modulated states of individual protein from our immune complexes, as manifested through global structural parameters. We analyzed the Root Mean Square Deviation (RMSD) and Radii of gyration (Rg) at different temperatures of each of the proteins under investigation. Calculating RMSD across the simulation time of protein backbones permits the quantification of the degree of conformational changes, as these changes may occur during MD simulations and can inform on the stability of proteins under different temperatures [1]. An overview of RMSD and *R*g curves is shown below.

**Supplementary Figure 2:**
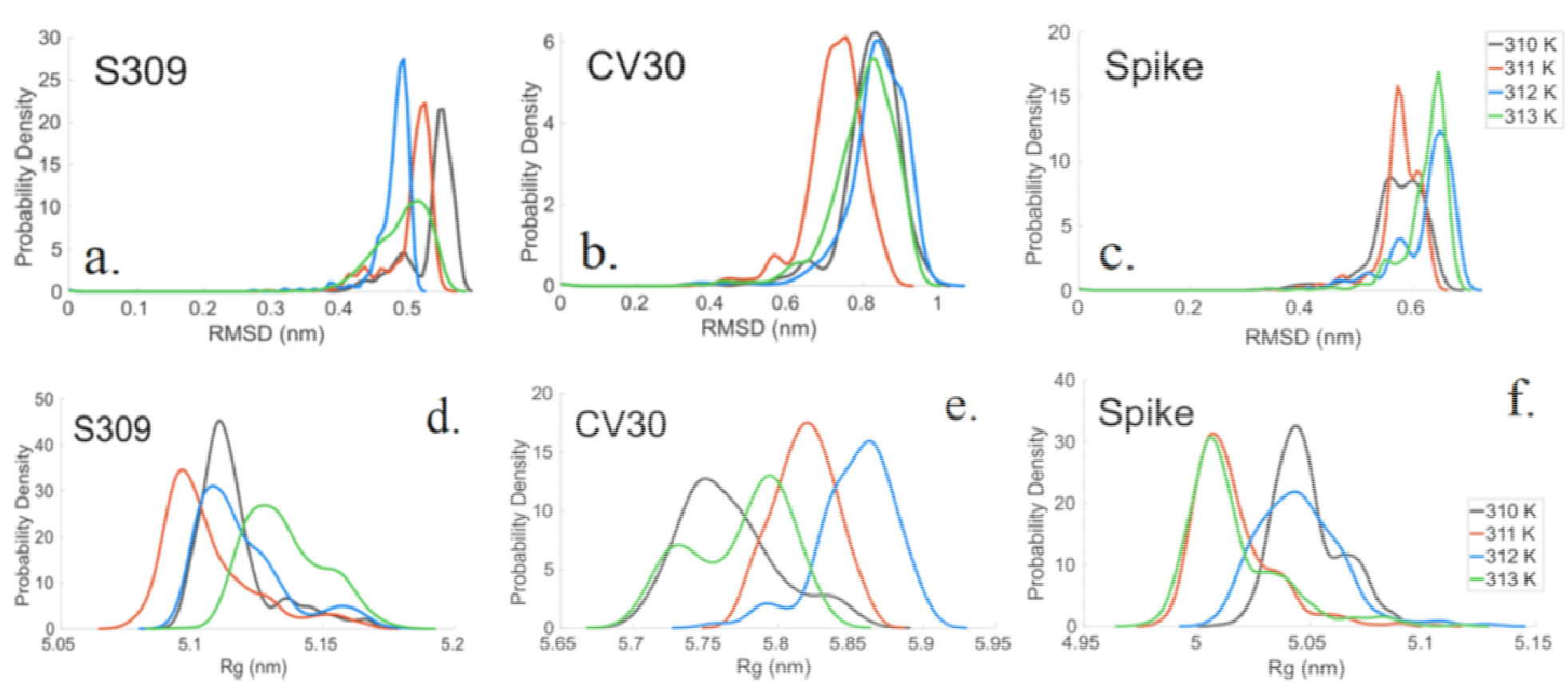
Upper panel: RMSD probability distribution function of the backbone C_α_ of S309 (a), CV30 (b) and viral Spike protein (c). Lower panel: *R*g probability distribution of S309 (d), CV30 (e) and viral Spike (f) at physiological and different fever temperatures.

For the S309, as well as for CV30, the least stable conformations are at 310K (average RMSD of 5.5 Å and 8.6 Å, respectively), whereas the most stable conformer is found at 312K (average of 5 Å) and at 311K (average of 7.5 Å), respectively (Supplementary Table 3). In contrast, it is notable that for the Spike, the backbone RMSD is more stable at 310K and 311K (average of 6 Å), while being less stable at 312K with an average of 6.8 Å. RMSD values we observed imply significant conformational changes at different febrile temperatures, such that Spikes become less stable with temperature increase, whereas both mAb become more stable at higher febrile temperatures.

We used *R*g to relate the compactness of proteins with changes in temperature during 400 ns of the simulations (Figure 2, lower panel, and Supplementary Table 3). *R*g monitors the dimensions of proteins during the simulations and correlates with the rate of folding and the maintenance of protein folds, with lowest *R*g indicating a tighter packing [2]. For S309, the values of *R*g converge at 310K-312K towards a similar value of around 5.1 nm; at around 200 ns, the most compact conformation is present at 311K (*R*g = 5.08. Importantly, at 313K, the least compact conformation is encountered (*R*g = 5.11), a situation that is opposite to CV30, where at the same temperature, the most compact conformation is found, whereas *R*g data at 311K present the highest increase. In the case of Spike, after a period of equilibration of around 50 ns, the most compact conformation is at 311K and 313K, whereas the least compact conformation is encountered at 310K. Interestingly, an increase in *R*g is observed around 150 ns and again around 225 ns, reflecting possible partial protein unfolding [3].

**Supplementary Figure 3.**
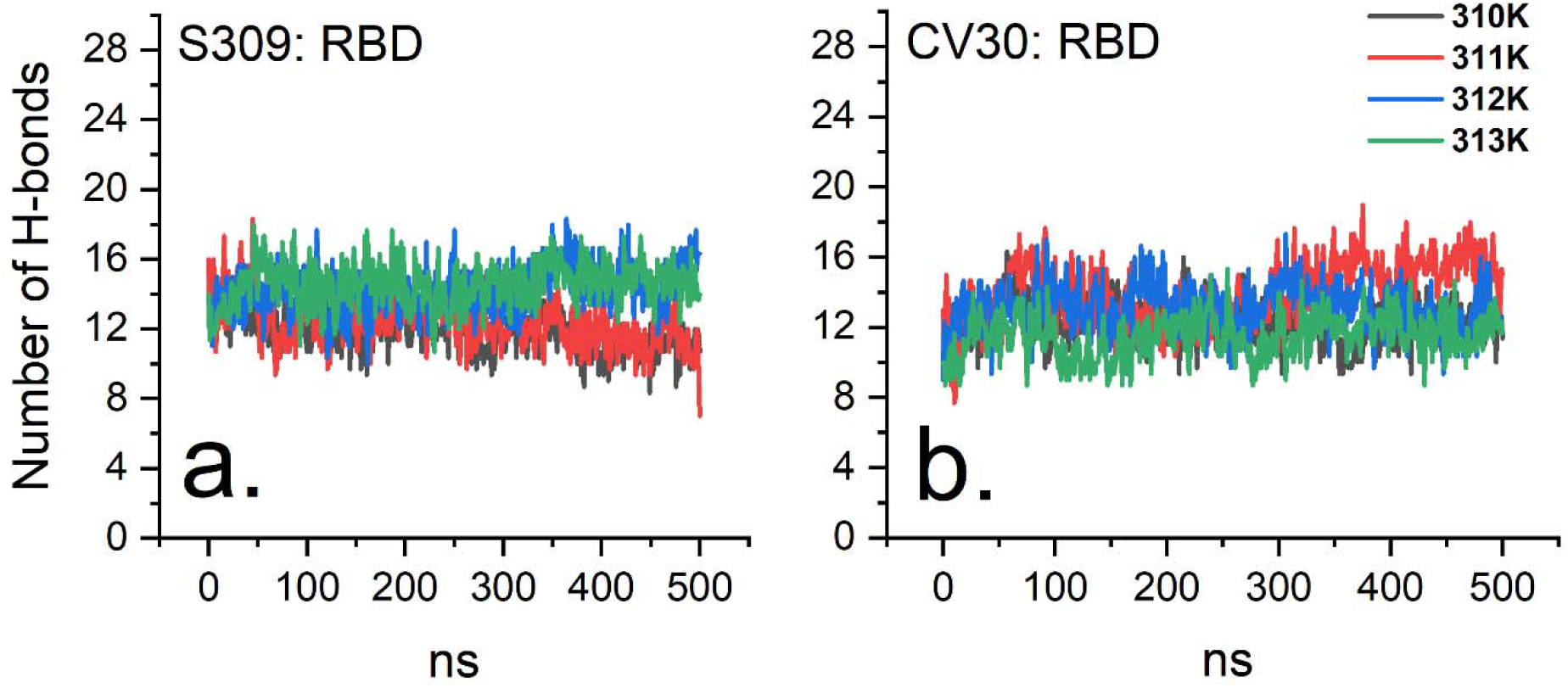
Evolution of hydrogen bond formation between the epitopes and the paratopes of the two immune complexes, under different temperature conditions.

