## Supplementary Table 1 for "Fever temperatures modulate intraprotein dynamics and enhance the binding affinity between monoclonal antibodies and the Spike protein from SARS-CoV-2"

|  | Chain A to B | Chain B to C | Chain A to C |
| --- | --- | --- | --- |
| S309, 310 K, 0 ns | 50.4 | 34.5 | 39.2 |
| S309, 310 K, 250 ns | 48.4 | 32.9 | 35.3 |
| S309, 310 K, 500 ns | 48.6 | 32.7 | 35 |
| S309, 311 K, 0 ns | 50.4 | 34.5 | 39.2 |
| S309, 311 K, 250 ns | 45.0 | 32.4 | 34.1 |
| S309, 311 K, 500 ns | 45 | 32.3 | 34.3 |
| S309, 312 K, 0 ns | 50.4 | 34.5 | 39.2 |
| S309, 312 K, 250 ns | 38.5 | 30.6 | 34.4 |
| S309, 312 K, 500 ns | 38 | 31.5 | 32.9 |
| S309, 313 K, 0 ns | 50.4 | 34.5 | 39.2 |
| S309, 313 K, 250 ns | 40.8 | 32.5 | 33.6 |
| S309, 313 K, 500 ns | 34.8 | 31.8 | 33.2 |

**Supplementary Table 1.** Changes in COM (Center of mass) between Spike RBDs at different temperatures and at different times during the simulations. Units are in Angstroms.

|  | Chain A to B | Chain B to C | Chain A to C |
| --- | --- | --- | --- |
| CV30, 310 K, 0 ns | 50.2 | 33.4 | 40.4 |
| CV30, 310 K, 250 ns | 45.6 | 33 | 36.2 |
| CV30, 310 K, 500 ns | 49.7 | 33 | 21.1 |
| CV30, 311 K, 0 ns | 49.3 | 35.2 | 38.9 |
| CV30, 311 K, 250 ns | 51.1 | 34.1 | 40.3 |
| CV30, 311 K, 500 ns | 53.6 | 33.7 | 39.2 |
| CV30, 312 K, 0 ns | 48.5 | 33.4 | 38.6 |
| CV30, 312 K, 250 ns | 58.1 | 34.3 | 40.9 |
| CV30, 312 K, 500 ns | 55.9 | 33.8 | 41.2 |
| CV30, 313 K, 0 ns | 48.8 | 33.7 | 39.7 |
| CV30, 313 K, 250 ns | 54.5 | 35 | 39.1 |
| CV30, 313 K, 500 ns | 51.5 | 35.2 | 34.9 |
