## Supplementary Table 2 for "Fever temperatures modulate intraprotein dynamics and enhance the binding affinity between monoclonal antibodies and the Spike protein from SARS-CoV-2"

**Supplementary Table 2.** Changes in BSA between Spike monomers throughout the simulations at the indicated temperatures. In brackets, percentage increase or decrease against values at 0 ns.

| BSA (Å^2^) | 310K CV30:Spike | 311K  CV30:Spike | 312K  CV30:Spike | 313K  CV30:Spike | 310K S309:Spike | 311K S309:Spike | 312K  S309:Spike | 313K  S309:Spike |
| --- | --- | --- | --- | --- | --- | --- | --- | --- |
| 0 ns | 20737 | 20593 | 20611 | 20673 | 19079 | 19093 | 19148 | 19017 |
| 500 ns | 21624 (+4%) | 21506 (+4%) | 22265 (+7%) | 22141 (+7%) | 21433 (+12%) | 21793 (+14%) | 23251 (+21%) | 21558 (+13%) |
