## Supplementary Table 3 for "Fever temperatures modulate intraprotein dynamics and enhance the binding affinity between monoclonal antibodies and the Spike protein from SARS-CoV-2"

**Supplementary Table 3.** Kolmogorov–Smirnov (KS) tests for RMSD and *R*g of individual proteins (mAb and Spike). Difference with statistical significance are indicated for all sets between the reference temperature (310K) and febrile temperatures (311K-313K). Values obtained using SciPy.

| KS test (RMSD) | S309 | CV30 | Spike |
| --- | --- | --- | --- |
| 310K – 311K | 4.39e-95 | 1.91e-84 | 1.34e-05 |
| 310K – 312K | 2.84e-143 | 1.50e-03 | 6.13e-58 |
| 310K – 313K | 7.63e-84 | 1.92e-04 | 3.85e-53 |
| 311K – 312K | 5.70e-108 | 2.27e-98 | 9.31e-71 |
| 311K – 313K | 6.78e-19 | 1.29e-50 | 1.06e-67 |
| 312K – 313K | 2.56e-35 | 1.12e-09 | 9.77e-05 |

| KS test (*R*g) | S309 | CV30 | Spike |
| --- | --- | --- | --- |
| 310K – 311K | 2.73e-46 | 9.05e-24 | 1.85e-118 |
| 310K – 312K | 8.33e-05 | 4.04e-35 | 1.34e-05 |
| 310K – 313K | 8.64e-94 | 4.12e-04 | 2.59e-107 |
| 311K – 312K | 2.16e-43 | 4.53e-17 | 1.47e-84 |
| 311K – 313K | 4.47e-129 | 2.71e-18 | 2.12e-01 |
| 312K – 313K | 6.41e-62 | 1.22e-45 | 1.73e-82 |
