## Supplementary Table 4 for "Fever temperatures modulate intraprotein dynamics and enhance the binding affinity between monoclonal antibodies and the Spike protein from SARS-CoV-2"

We determined the extent of hydrogen bonding between RBDs and mAb. We investigated the dynamics of hydrogen bond formation between the epitopes and the paratopes of our complexes, under relevant temperature conditions.

**Supplementary Table 4. Total counts of hydrogen bonds between RBD_1_ (residues 330-529) and the two paratopes under study, at physiological and at febrile temperatures. Simulation time is 500 ns. In brackets, percentage increase or decrease in the counts of hydrogen bonds against the reference values at 310K.**

| Number of H-bonds | 310K | 311K | 312K | 313K |
| --- | --- | --- | --- | --- |
| CV30: RBD_1_ | 6135 | 6995 (+14%) | 6549 (+6%) | 5751 (- 6%) |
| S309: RBD_1_ | 5952 | 6188 (+3%) | 7149 (+17%) | 7270 (+19%) |

A common feature of the two systems was the increase in H-bonds at febrile temperatures, peaking at 311K for the CV30:RBD system (14% increase compared to values at 310K). A corresponding increase in stability of the S309:RBD system is observed across all febrile temperatures, peaking at 312K and at 313K (a 19% average increase versus values obtained at physiological temperature).
